# Depletion of Protein Phosphatase 1 Results in Persistent Activation of the Integrated Stress Response and Age-Associated Exacerbation of Pulmonary Veno-Occlusive Disease

**DOI:** 10.1101/2024.04.16.589837

**Authors:** Amit Prabhakar, Meetu Wadhwa, Prajakta Ghatpande, Rahul Kumar, Brian B. Graham, Giorgio Lagna, Akiko Hata

**Affiliations:** Cardiovascular Research Institute, University of California, San Francisco, San Francisco, CA 94143, USA; Department of Anesthesia and Perioperative Care, University of California, San Francisco, San Francisco, CA 94143, USA; Department of Radiology, University of California, San Francisco, San Francisco, CA 94143, USA; Lung Biology Center, Pulmonary and Critical Care Medicine, Zuckerberg San Francisco General Hospital, CA, 94110, USA; Department of Biochemistry and Biophysics, University of California, San Francisco, San Francisco, CA 94143, USA

## Abstract

Pulmonary veno-occlusive disease (PVOD) is a form of pulmonary hypertension that affects individuals across the age spectrum. PVOD is characterized by the obstruction of small pulmonary vessels, causing increased pulmonary artery (PA) pressure and leading to right ventricular heart (RV) failure. Previous research showed that the administration of Mitomycin-C (MMC) in rats mediates PVOD through the activation of the eukaryotic initiation factor 2 (eIF2) kinase PKR and the integrated stress response (ISR), resulting in the impairment of vascular endothelial junctional structure and barrier function. In this study, we reveal that older rats experience more severe pulmonary vascular remodeling and RV hypertrophy than younger rats after MMC treatment due to lower levels of protein phosphatase 1, leading to prolonged eIF2 phosphorylation and ISR activation. We demonstrate that pharmacological blocking of the PKR-ISR pathway mitigates PVOD symptoms in both age groups, suggesting targeting the PKR-ISR axis as a potential PVOD therapeutic strategy.

## Introduction

PVOD/pulmonary capillary hemangiomatosis is a subclass of pulmonary hypertension (PH) characterized by the progressive remodeling and obstruction of small PAs, pulmonary veins (PVs), and capillaries (PCs), leading to increased PA pressure and RV failure (1, 2). PVOD can begin at any age and affects both males and females (1). Because there is no effective therapy for PVOD, a mortality rate is 72% within 1 year of diagnosis(1). The vascular lesions in human PVOD patients also include thrombosis, fibrous intimal proliferation, and dilatation and proliferation of PCs(1). Biallelic loss-of-function/expression mutations in the *EIF2AK4* gene, which encodes the general control nonderepressible 2 (GCN2), one of four kinases that phosphorylate the eukaryotic translation initiation factor 2 (eIF2) and regulate cap-dependent translation, are the primary genetic cause of PVOD(3, 4). PVOD is often misdiagnosed as pulmonary arterial hypertension (PAH) due to similarities in radiographic findings and the overlap of gene mutations between PVOD and PAH(3, 4). Pharmacologically, the administration of chemotherapeutic agents, such as mitomycin C (MMC), bleomycin, and cisplatin, has been implicated in the onset of PVOD (5–7). Specifically, MMC administration to rats induces a spectrum of PVOD-like phenotypes, including right ventricular (RV) hypertrophy and changes in the pulmonary vasculature, including medial thickening and obstruction of the lumen in PAs and PVs, fibrous growth in the intima and adventitia, and thrombosis (6, 8–11). Therefore, the MMC-mediated PVOD rat model offers valuable insights into the pathogenesis of PVOD and provides a platform for testing potential therapeutic interventions.

The pulmonary vascular endothelium (also known as intima) is composed of a monolayer of endothelial cells that form the lining of PAs, PVs, and PCs(12). VE-Cad is a critical component of the adherens junctions (AJs) in endothelial cells, playing an essential role in maintaining permeability and vascular integrity(12). Previous studies have demonstrated that VE-Cad localizes at the AJs in complex with Rad51 (hereafter referred to as VE-Cad:Rad51 complex or VRC) (11). The interaction between Rad51 and VE-Cad is essential for the stability of VE-Cad, as well as for preserving the integrity of endothelial cell-cell junctions and barrier function(11). MMC treatment leads to the depletion of VE-Cad and Rad51 in endothelial cells partially through the ubiquitin-proteasome-dependent degradation of Rad51, which results in the destabilization of VE-Cad(11). Additionally, MMC treatment contributes to VE-Cad depletion by promoting the release of the VRC from endothelial cells into the extracellular space(11). Inhibiting Rad51 degradation by silencing of the E3 ubiquitin ligase for Rad51 Fbh1 or excess amount of Rad51 in the cell protects VE-Cad after MMC treatment and maintains the barrier function(11). Therefore, the depletion of the VRC following MMC treatment is likely associated with pulmonary vascular remodeling in PVOD patients are all affected.

The integrated stress response (ISR) is an evolutionary conserved adaptive intracellular mechanism involved in the maintenance of homeostasis upon changes in the cellular environment and pathological stimuli(13). However, the activation of ISR is also associated with various disorders and age-related ailments in human(13). The ISR is initiated by the activation of stress-responsive kinases, which include GCN2, double-stranded RNA-activated protein kinase (PKR), PKR-like endoplasmic reticulum kinase (PERK) and heme-regulated inhibitor kinase (HRI). Upon activation, these kinases phosphorylate serine-51 (Ser^51^) of the alpha subunit of eukaryotic initiation factor 2 (eIF2α)(14). When eIF2α is phosphorylated at Ser^51^ (p-eIF2α), it associates with eIF2B, a guanine nucleotide exchange factor for eIF2, and cap-dependent translation is attenuated(13). Under the condition which general protein synthesis is suppressed, translation of specific transcripts, such as *cyclic AMP-dependent transcription factor 4 (ATF4)*, a master transcriptional of stress genes, is promoted and ATF4-mediated gene regulation orchestrates ISR signaling pathway (15). Moreover, the ATF4 mRNA, which bears upstream open reading frames (uORFs), is preferentially translated when general translation is suppressed (16). We found that when rats are administered with MMC, the ISR pathway is activated by PKR, resulting in the impairment of vascular endothelial homeostasis and barrier function(11). The treatment with the antagonist of PKR or ISR attenuates the development of PVOD phenotypes mediated by MMC(11).

When the cellular environment has been changed and the environmental stressor is removed, the ISR signal is terminated via the dephosphorylation of p-eIF2α at Ser^51^, which then leads to the restoration of global translation and other physiological activities(13). Dephosphorylation of eIF2α(Ser^51^) is mediated by protein phosphatase 1 (PP1) that is composed of a catalytic subunit PP1c (also known as PPP1C) and either stress-induced regulatory subunit growth arrest and DNA damage-inducible protein (GADD34, also known as PPP1R15A) or constitutive-expressed regulatory subunit constitutive repressor of eIF2α phosphorylation (CReP, also known as PPP1R15B) (13, 17). Unlike CReP, which serves as a regulator of PP1c in unstressed cells, GADD34 is induced by the activation of the ISR under stress. This induction occurs through the transcriptional activation of the GADD34 gene by ATF4(18), in combination with the preferential translation of the GADD34 transcripts under the ISR activation and global translational inhibition because, like the ATF4 transcripts, they contain uORFs (19). When PP1 (PP1c: GADD34) is formed, it eIF2α dephosphorylation of eIF2α at Ser^51^ and terminates ISR. Thus, PP1 (PP1c: GADD34) provides a critical negative feedback loop to restore protein synthesis once the ISR pathway is activated.

In this study, we report that old rats exhibit an augmented ISR activity compared to young rats. Upon MMC treatment, old rats develop more severe PVOD phenotypes than young rats. We found that treatment with MMC leads to the depletion of both PP1c and GADD34 proteins, which results in prolonged phosphorylation of eIF2 and activation of the ISR. Administering the PKR antagonist C16 or the ISR inhibitor ISRIB after the onset of PVOD phenotypes effectively reversed the depletion of the PP1c:GADD34 complex and ameliorated the cardiopulmonary phenotypes of PVOD in both young and old rats. These findings suggest that pharmacological intervention targeting the PKR-ISR axis is an effective strategy for mitigating PVOD, regardless of the animal’s age.

## Results

### Increased ISR activity in old rats

ISR activity has been linked to the aging process(20) and the inhibition of ISR by ISRIB has been found to mitigate the effects of aging(21). We examined the ISR activity in the lung tissues and compared it between young (9-10 weeks old) and old (1.2-1.5 year old) Sprague Dawley rats (**Fig. 1A**). The amount of active form of PKR (p-PKR) relative to total PKR (p-PKR/t-PKR), p-eIF2α relative to total eIF2α (p-eIF2α/t-eIF2α), and cyclic AMP-dependent transcription factor 4 (ATF4), a master transcriptional regulator for activating the ISR pathway(15), were all elevated in old rats compared to young rats, 1.2-fold, 2-fold, and 1.5-fold, respectively (**Fig. 1A and Supple. Fig. S1**). Moreover, the mRNA levels of the genes induced upon activation of the ATF4-ISR pathway, such as *ATF3*, *ATF4*, and *PKR,* were 1.2-fold, 2-fold, and 14-fold higher in old rats than young rats, respectively (**Fig. 1B**). The amount of another eIF2α kinase GCN2 detected in the lung was comparable between young and old rats (**Fig. 1A**). The levels of VE-Cad and Rad51 were 40% and 39% lower in old rats than young rats, respectively (**Fig. 1A**). There was no age-related change in the amount of GCN2 (**Fig. 1A**).

**Fig.1.**
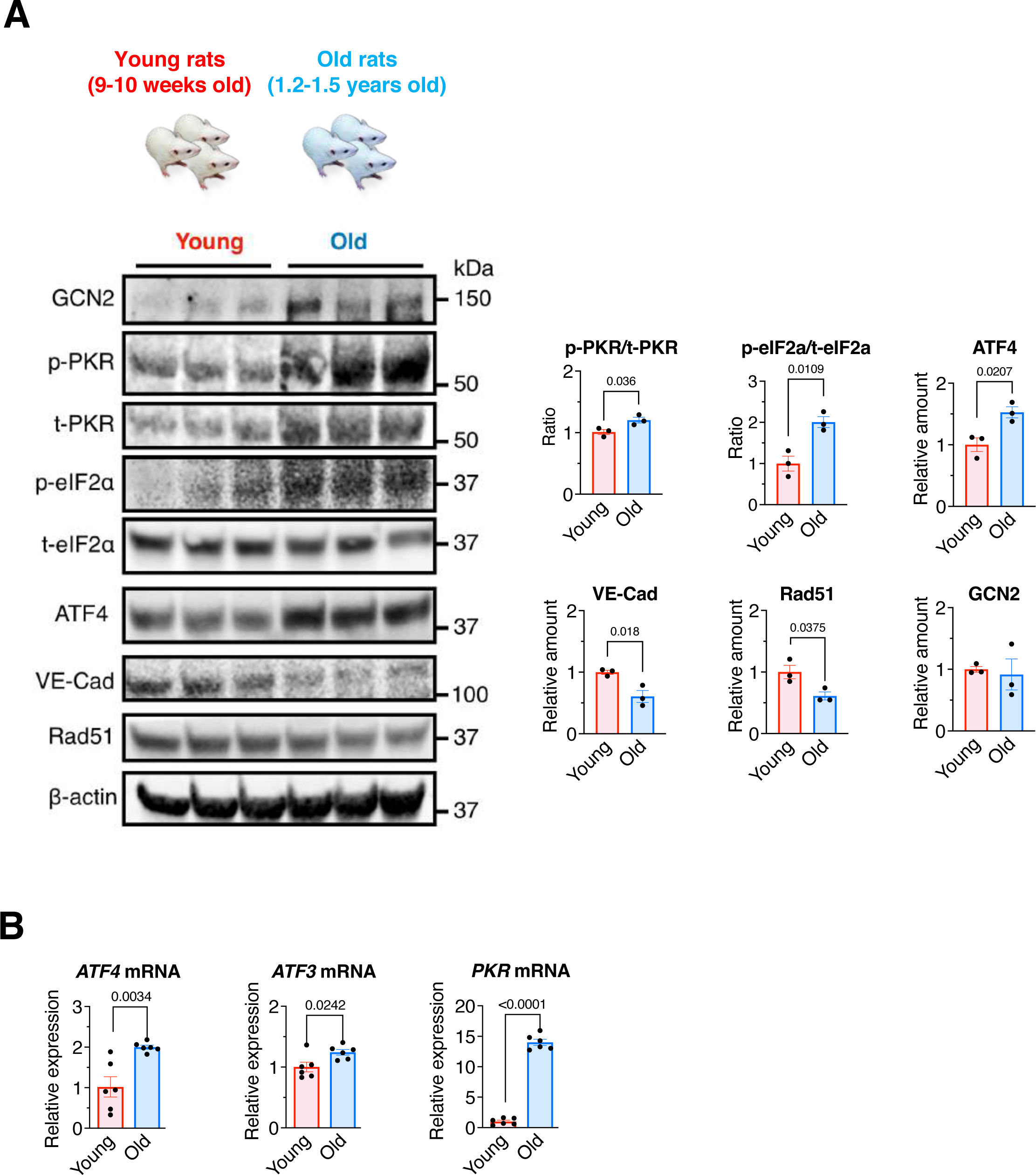
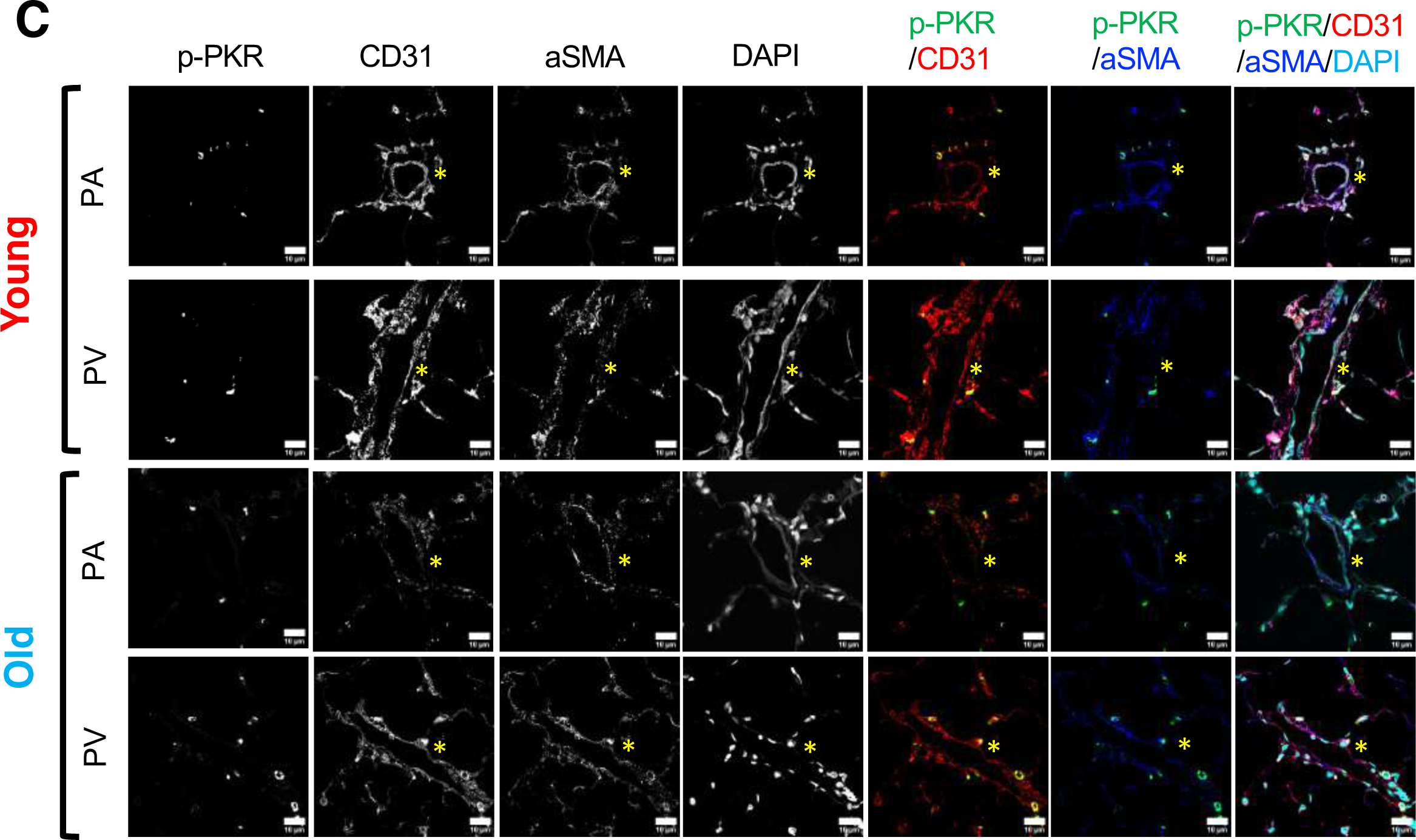
Age-dependent increase in the basal ISR activity A. Lung lysates of young and old rats were subjected to immunoblot of the indicated proteins (left). The amount of indicated proteins normalized to β-actin, the ratio of p-PKR/t-PKR and p-eIF2α/t-eIF2α are shown as mean±SEM (right). n=3 independent samples per condition. **B.** The level of mRNAs of ATF4 target genes, such as ATF4, ATF3, and PKR, in the lung young and old rats were analyzed by qRT-PCR and shown as mean±SEM after normalized to the GAPDH mRNA. n=3 independent samples. **C.** Lung lysates from young and old rats were subjected to IF staining with anti-p-PKR (green), anti-CD31 (red), and anti-aSMA (blue) antibody and the images of PAs and PVs are shown. Cell nuclei were stained with DAPI (cyan). Asterisks indicate the location of PAs and PVs. Scale bar=10’m. Statistical analysis was performed using two-tailed Student’s t-test with p<0.05.

Immunofluorescence (IF) staining of p-PKR in the lung sections detected significantly higher p-PKR signal compared to young rats in CD31-positive pulmonary vascular endothelial cells but not in smooth muscle cells that are positive with the α-smooth muscle actin (aSMA) (**Fig. 1C**). These results demonstrate an age-associated elevation of the basal ISR activity in the lung as a result of PKR activation.

### Reversal of PVOD phenotypes in young and old rats

Because sporadic PVOD case are predominantly individuals with older age(22), we compared MMC-induced PVOD phenotypes between young and old rats. Twenty-four days after the administration of MMC, rats were treated with vehicle, the PKR antagonist C16 (23), or the ISR inhibitor ISRIB (24, 25) for 8 days, followed by the analysis of cardiovascular phenotypes (**Fig. 2A**). The rats treated with MMC but not with C16 nor ISRIB (MMC/vehicle) exhibited the hallmarks of pulmonary hypertension phenotype, such as elevation of PA pressure shown as right ventricular systolic pressure (RVSP; from 25.0 mmHg to 34.1 mmHg in young rats and from 24.4 mmHg to 34.6 mmHg in old rats) and RV hypertrophy shown as right ventricle RV/LV + septum (S) weight ratio (RV/LV+S; from 0.21 to 0.31 in young rats and from 0.21 to 0.34 in old rats) (**Fig. 2A**). The old rats demonstrated slightly higher levels of RVSP and RV/LV+S ratio than young rats (**Fig. 2A**). When treated with either C16 or ISRIB, RVSP and RV/LV+S ratio were reversed to the levels similar to control (vehicle/vehicle-treated) young and old rats (**Fig. 2A**). H&E staining of the lung showed MMC/vehicle-treated both young and old rats developed significant medial hyperplasia/hypertrophy in pulmonary arterioles (PAs) and muscularization of pulmonary venules (PVs)(6, 8) (**Fig. 2B**). Both microvessels (<50 μm in diameter) and medium-sized vessels (50-80 μm in diameter) underwent medial thickening, damaged intima resulting in the occlusion of vessels, in young and old rats following MMC treatment (**Fig. 2B**). The fraction of vessels with moderate (4.07% to 35.56% increase in young; 7.78% to 41.85% increase in old rats) to severe (0.76% to 29.38% increase in young; 1.45% to 30.51% increase in old rats) medial thickening elevated from 4.83% to 64.90% after MMC treatment in young rats and from 9.23% to 72.36% in old rats, resulting in narrowing or complete obstruction of the lumen (**Fig. 2B**). Thus, old rats develop more severe vascular remodeling than young rats post-MMC. When rats were treated with C16 or ISRIB, pulmonary vascular remodeling in both young and old rats were reversed and indistinguishable from MMC-untreated rats (**Fig. 2B**). Trichrome staining indicated that fibrous growth of intima and adventitia in young and old rat post-MMC, which was mitigated by C16 or ISRIB treatment in both young and old rats (**Fig. 2C**). The lung lysates of MMC-treated young and old rats exhibited higher levels of in p-PKR/t-PKR ratio, p-eIF2α/t-eIF2α ratio, ATF4 than untreated rats, demonstrating the ISR activation by MMC (**Fig. 2D**). The levels of ISR activity in old rats were higher than in young rats (**Fig. 2D**). Delayed C16 or ISRIB treatment after the development of MMC-induced cardiovascular phenotypes in young and old rats reversed to the levels like control (vehicle/vehicle-treated) rats (**Fig. 2E**). Furthermore, delayed C16 or ISRIB treatment reversed the amount of VE-Cad and Rad51, which were depleted from vascular endothelial cells after MMC treatment (**Fig. 2E**). The transcriptional activation of ATF4 target genes (*ATF3, ATF4*, and *PKR*) post-MMC treatment was reversed by either C16 or ISRIB treatment in young and old rats, confirming the inhibition of the PKR-ISR axis by C16 and ISRIB (**Fig. 2F**). These results confirm that a pharmacological intervention of the PKR-ISR axis by C16 or ISRIB can reverse MMC-mediated PVOD phenotypes even after they were fully developed.

**Fig. 2.**
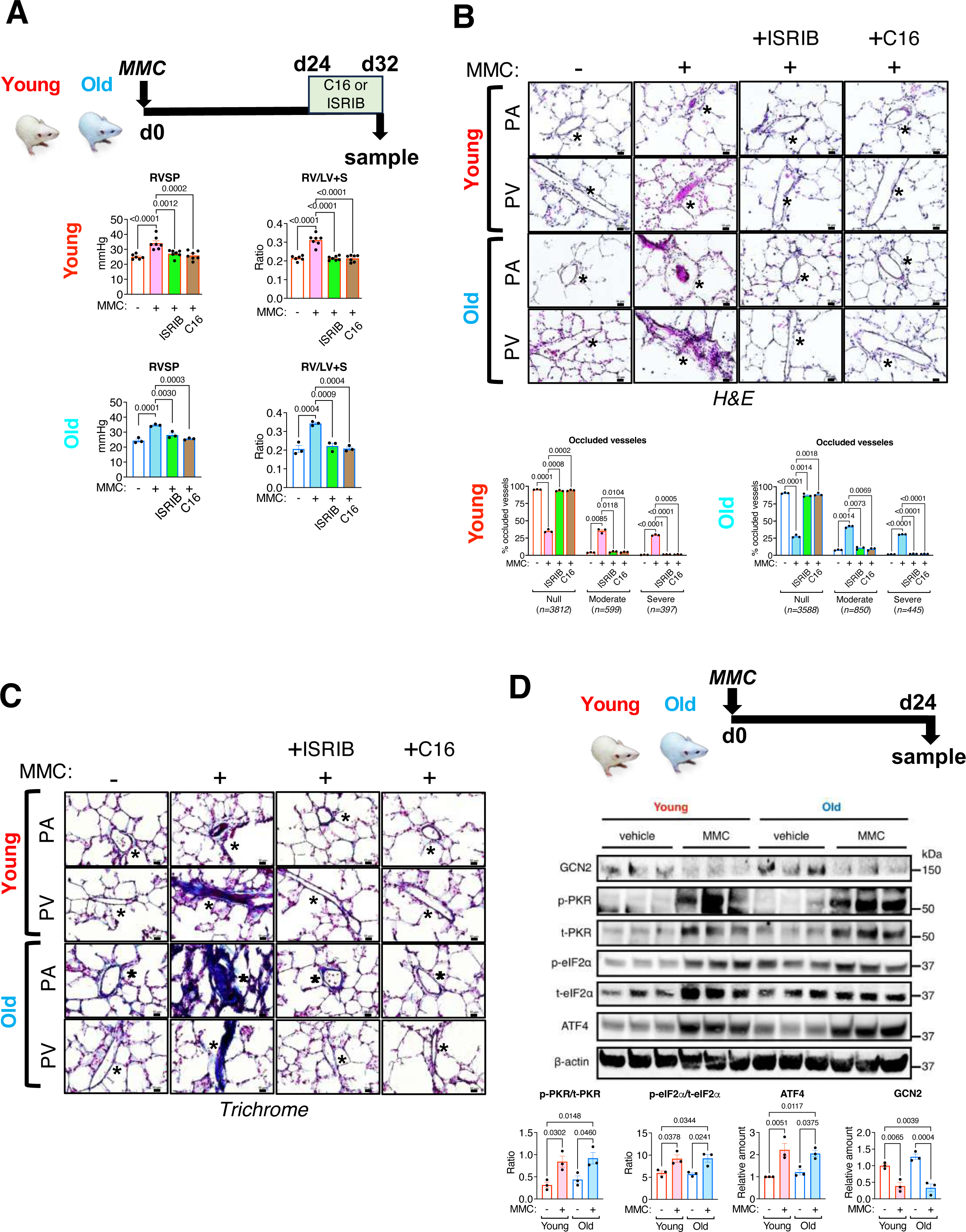
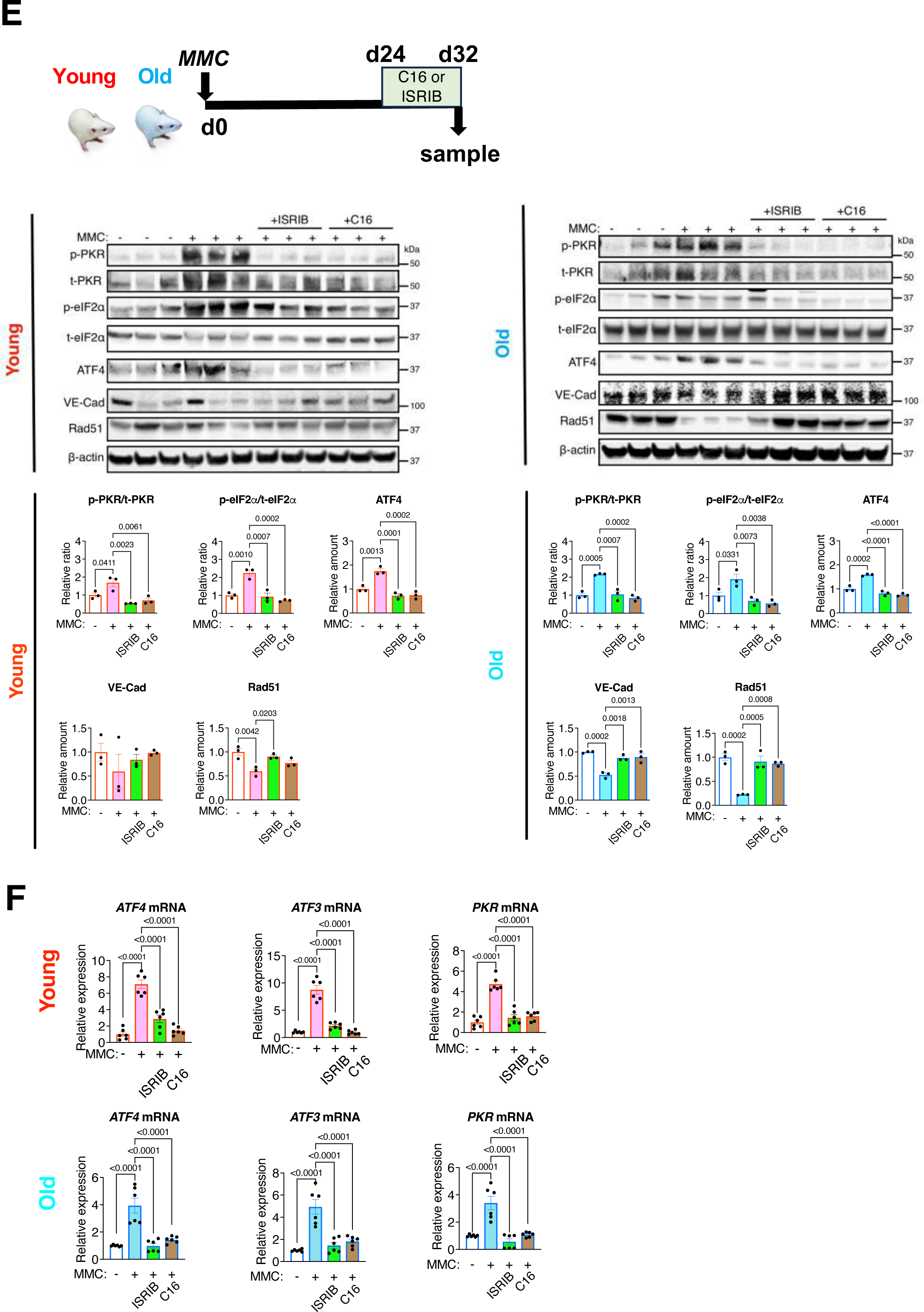
Administration of C16 or ISRIB 24 days after MMC treatment reverses PVOD phenotypes A. A scheme of delayed C16 and ISRIB treatment on d24-32 after the administration of MMC on d0 in old and young rats (top). RVSP and RV/LV+S ratio in vehicle-or MMC-exposed rats with or without C16 or ISRIB treatment are shown as mean±SEM (bottom). n=3-7 independent samples. **B.** H&E staining of pulmonary vasculature (Pas and PVs) in young and old rats treated with vehicle or MMC with or without C16 or ISRIB treatment was performed (top). Total number of microvessels were analyzed, distributed in three categories based on medial thickening/occlusion of pulmonary vasculature as null (no occlusion), moderate (25-50% occlusion), and severe (50-100%) in young (bottom, left) and old (bottom, right) rats. Data is presented as a faction (%) of vessels in each category relative to total vessels count and shown as mean±SEM. Scale bar=10 mm. n=3 independent experiments. **C.** Trichrome staining of pulmonary vasculature (PAs and PVs) in young and old rats treated with vehicle or MMC with or without delayed C16 or ISRIB treatment was performed on d32 and shown (left). Trichrome stain intensity was quantitated, converted to the value relative to the intensity of vehicle-treated young rats, and shown as mean±SEM (right). Scale bar=10 mm. n=3 independent experiments. **D.** A scheme of MMC treatment in young and old rats (top). Lung lysates of young and old rats 24 days after vehicle or MMC administration were subjected to immunoblot of the indicated proteins (left). The amount of indicated proteins normalized to β-actin, the ratio of p-PKR/t-PKR and p-eIF2α/t-eIF2α are shown as mean**±**SEM (right). n=3 independent samples per condition. **E.** Immunoblot analysis of indicated proteins in total lung lysates from vehicle (-), MMC (+) with or without C16 or ISRIB in young (top, left) and old rats (top, right). The amount of indicated proteins relative to β-actin are shown as mean**±**SEM (bottom). n=3 independent samples. **F.** The levels of mRNAs of ATF4 target genes, such as ATF4, ATF3, and PKR, in the lung of from young and old rats administered with vehicle, MMC, MMC+ISRIB, or MMC+C16 were analyzed by qRT-PCR and shown as mean**±**SEM after normalized to the GAPDH mRNA. n=3 independent samples. Statistical analysis was performed using one-way ANOVA with Tukey’s multiple comparisons test or two-way ANOVA with Tukey’s multiple comparisons test with *p*<0.05.

### Reduction of the intracellular VRC amount is reversed by the inhibition of PKR or ISR

We previously found that upon MMC treatment, the VE-Cad:Rad51 complex (VRC) is released from the vascular endothelium into the circulation upon MMC treatment in rats, which results in the depletion of VRC in the vascular endothelium. MMC-treated young and old rats exhibited increased levels of VRC in the plasma (**Fig. 3, Plasma**). Conversely, the VRC in the lung lysates was reduced (**Fig. 3, Lung**). C16 or ISRIB treatment blocked the release of the VRC into the circulation post-MMC treatment (**Fig. 3, Plasma**) and rescued the VRC in the lung lysates (**Fig. 3, Lung**). These results suggest that the inhibition of the PKR-ISR axis is effective in reversing PVOD phenotypes regardless of the animal’s age and restore endothelial cell-cell junctions and barrier function by restoring the VRC.

**Fig. 3.**
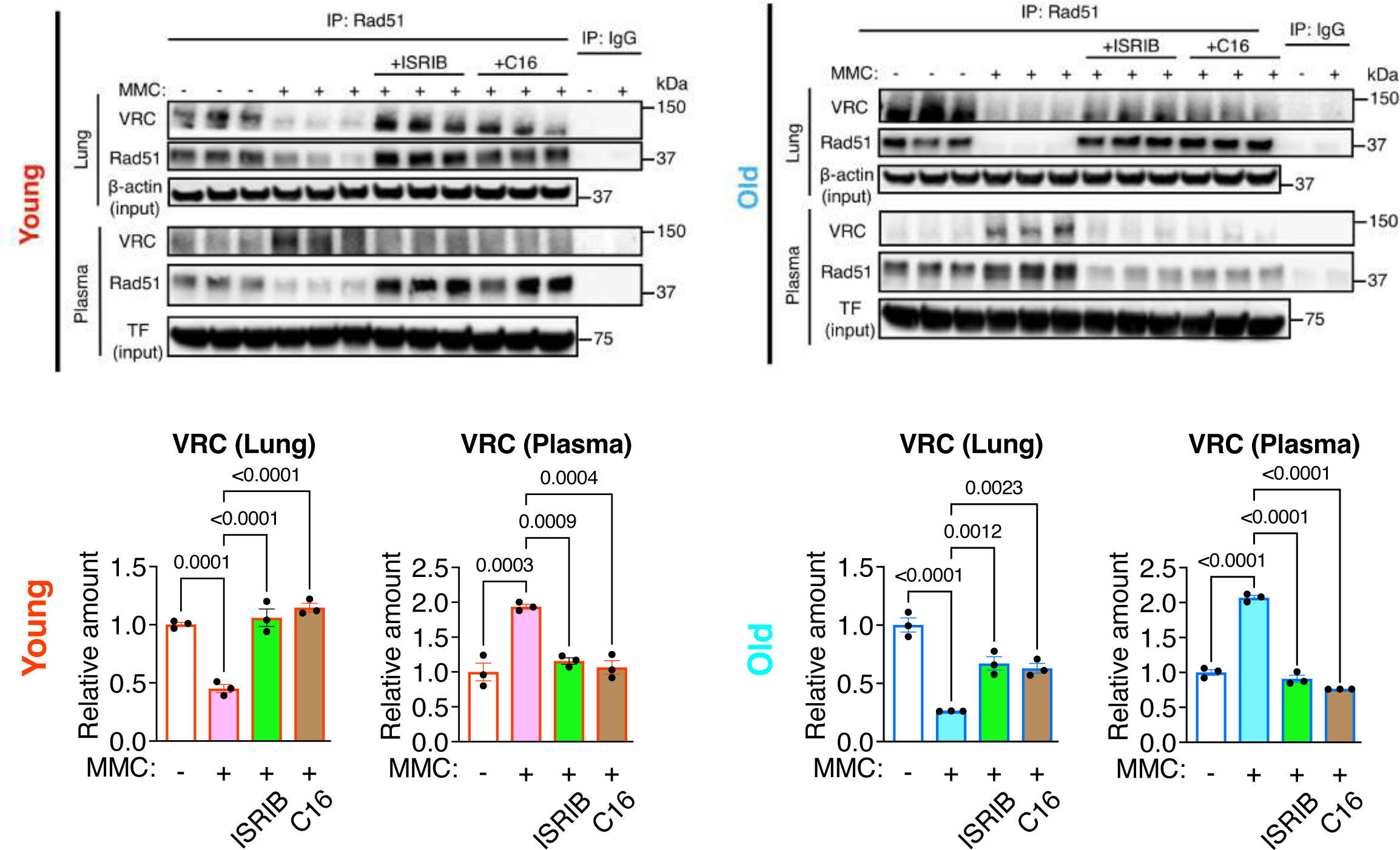
Inhibition the PKR-ISR axis blocks the accumulation of VRC in the circulation and restores it in vascular endothelial cells. Total lung lysates and plasma samples from vehicle (-), MMC (+), MMC+ISRIB, and MMC+C16 treated young (top, left) and old rats (top, right) were subjected to immunoprecipitation (IP) by an anti-Rad51 antibody or nonspecific IgG (control), followed by immunoblot analysis of VE-Cad (for VRC) and Rad51 (top). The total cell lysates and plasma samples without IP were subjected to immunoblot with anti-β-actin (for lung) and anti-transferrin (TF) antibody (for plasma) as loading control. The relative amount of VRC is shown as mean**±**SEM (bottom). n=3 independent samples. Statistical analysis was performed using one-way ANOVA with Tukey’s multiple comparisons test with *p*<0.05.

### The lack of GADD34 induction and constitutive ISR activation following MMC treatment

Dephosphorylation of eIF2α by PP1 (PP1c:GADD34 complex) is critical in terminating the ISR signals and restoring protein synthesis after cellular stress(13, 17). Therefore, we assessed the levels of the PP1c and GADD34 mRNA and protein in young and old rats. After MMC treatment, the levels of the PP1c mRNA were decreased 48% and 77% in young and old rats, respectively (**Fig. 4A**). Unlike PP1c mRNA, we observed a 9.0-fold and 5.9-fold increase in the GADD34 mRNA following MMC treatment in young and old rats, respectively (**Fig. 4A**). Moreover, the GADD34 mRNA levels returned to the baseline (vehicle-treated rats) after the treatment with ISRIB or C16 (**Fig. 4A**). These results confirm that MMC-mediated activation of the ISR leads to the transcriptional activation of the GADD34 gene by ATF4 in young and old rats as expected(13), which can be reversed by the inhibition of PKR or ISR by C16 or ISRIB. Despite a significant increase in the GADD34 mRNA levels, GADD34 protein amount decreased 51% and 49% after MMC treatment in young and old rats, respectively (**Fig. 4B**). Similarly, PP1c protein amount decreased 81% and 80% post-MMC treatment in young and old rats, respectively (**Fig. 4B**). Consequently, the amount of PP1 (PP1c:GADD34 complex) in MMC-treated young and old rats reduced to 41% and 26% of vehicle-treated rats, respectively (**Fig. 4b**). The reduction of PP1 complex amount by MMC was reversed upon ISRIB or C16 treatment (**Fig. 4B**). These results provide evidence that the depletion of PP1 by MMC results in the persistent eIF2 phosphorylation and maladaptive ISR activation, which in turn mediates pulmonary vascular endothelial dysfunction and vascular remodeling (**Fig. 4C**). C16 or ISRIB treatment restores PP1 in the vascular endothelium, terminates prolonged ISR activities, and ameliorates vascular remodeling in PVOD (**Fig. 4C**).

**Fig. 4.**
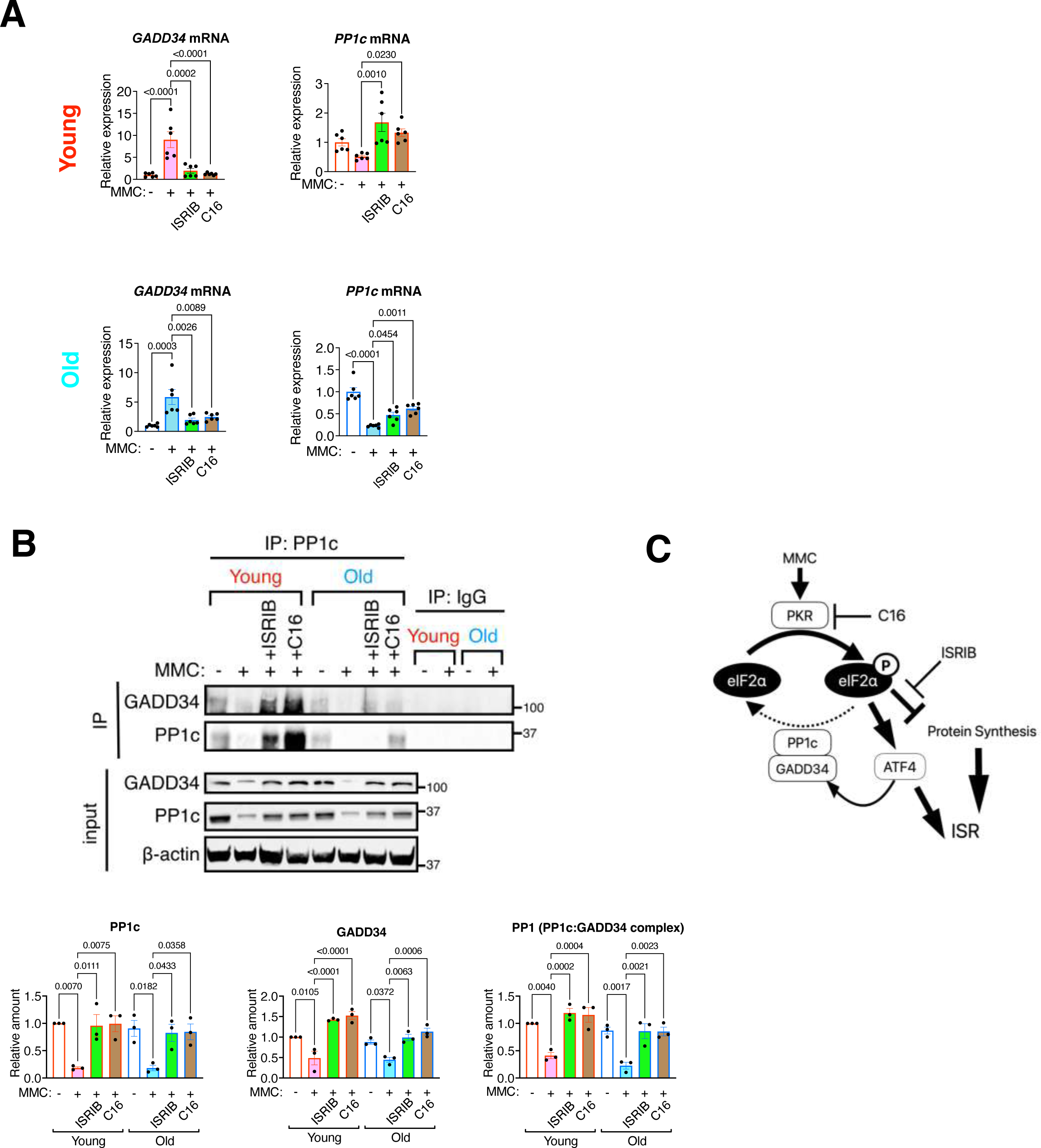
Reduced PP1 activity results in prolonged ISR activation A. The levels of GADD34 and PP1c mRNAs in the lung of young and old rats treated with vehicle (-), MMC (+), MMC+ISRIB, and MMC+C16 were analyzed by qRT-PCR, normalized to GAPDH, and shown as mean±SEM. n=3 independent samples. **B.** The lung lysates of levels of young and old rats treated with vehicle (-), MMC (+), MMC+ISRIB, and MMC+C16 were subjected to immunoprecipitation (IP) with an anti-PP1c antibody or nonspecific IgG (control), followed by immunoblot with anti-GADD34 antibody to assess the amount of the PP1c:GADD34 complex (left). Input samples were subjected to immunoblot analysis with PP1c, GADD34, and β-actin antibody (loading control). The amount of PP1c and GADD34 in input samples was normalized to β-actin and shown as mean±SEM (bottom left and middle). The amount of GADD34 in the IP samples was normalized to β-actin and shown as mean±SEM (bottom right). n=3 independent samples. Statistical analysis was performed using two-tailed Student’s t-test and one-way ANOVA with Tukey’s multiple comparisons test with p<0.05. **c.** Immunoblot of the GADD34 and PP1c proteins in the lung lysate from vehicle (-), MMC (+), MMC+ISRIB, and MMC+C16 treated young and old rats were normalized to β-actin and shown as mean±SEM. n=3 independent samples. **C.** The schematic depicts the pathway of constitutive ISR activation associated with aging, leading to exacerbated PVOD phenotypes. The reduction of PP1 impedes the dephosphorylation of eIF2α, resulting in a global translational inhibition within vascular endothelial cells. This inhibition contributes to age-related endothelial dysfunction and the development of PVOD. The antagonists of PKR (C16) and ISR (ISRIB) inhibit sustained ISR activation, potentially reversing PVOD phenotypes.

## Discussion

In this study, we demonstrate that the antagonist of PKR or ISR effectively reverses the phenotypes associated with PVOD, such as pulmonary vascular obstruction and RV hypertrophy, in both young and old rats. These results further confirm that the intervention of the PKR-ISR by C16 or ISRIB might be an effective treatment even for the advanced stage of PVOD. The ISR pathway is an evolutionarily conserved adaptive mechanism to a changing environment and maintain homeostasis and is fundamental to organismal resilience and longevity (13, 26). Maladaptive activation of ISR is linked to various age-associated diseases, such as Parkinson’s disease, Alzheimer’s disease, multiple sclerosis, and Amyotrophic lateral sclerosis, and the inhibition of ISR by ISRIB mitigates these conditions (13, 20, 26). Moreover, increased levels of p-eIF2α has been detected in various organs from old animals (20, 27). We found that the basal levels of ISR in old rats is higher than young rats. After MMC treatment, PVOD in old rats exhibited more severe form of PVOD than young rats, indicating the higher levels of the ISR activity contribute to more pronounced pathological changes in PVOD model rats. PVOD affects both females and males of all age-groups ranging from 8 weeks to the 7^th^ decade of life, but sporadic PVOD case are predominantly male with older age(22). Together with the results presented in this study, we speculate that exposure to multiple stress during aging and the prolonged activation of ISR and constitutive downregulation of translation over time result in the higher incidence of PVOD among older age individuals.

The eIF2 kinases constitute the first step of ISR regulation and several studies using rodents report age-related changes in their expression(20). A comparison of adult and old male mice demonstrated an increase of PKR protein levels in all tested tissues, including kidney, liver, colon, brain, testes, pancreas, lung, and heart(27). Human muscle biopsies from donors ranging from 20 to >80 years of age also show increasing PKR abundance with age(27, 28). Thus, it is plausible that the elevated amount of t-PKR in old rats is the result of normal aging process.

We find that the diminished levels of the catalytic subunit of PP1 (PP1c), coupled with the inability to induce the regulatory subunit of PP1 (GADD34) in the pulmonary vascular endothelium after MMC treatment, lead to the persistent activation of the ISR signals. This results in a prolonged inhibition of protein synthesis. Consequently, what begins as a transient, “adaptive” ISR signal converts into a constitutive, “maladaptive” signal, resulting in the pathological remodeling of the pulmonary vasculature. PP1 is known to play a critical role in regulating cognitive functions, such as learning and memory(29), synaptic transmission, and plasticity(30). Moreover, the decline in memory associated with aging has been attributed to the depletion of PP1c protein (31). Therefore, we speculate that the reduction in PP1 activity may underlie various age-associated pathologies, including PVOD and memory loss. Our study highlights the therapeutic potential of restoring PP1 as a treatment for PVOD, alongside the attenuation of the PKR-ISR pathway.

## Methods

Reagents, kits, antibodies, PCR primers, siRNAs, instruments, and software used in the study are listed in the Supplementary Information.

### Animal care and use

All animal experiments were conducted in accordance with the guidelines of the Institutional Animal Care and Use Committee (IACUC) of University of California, San Francisco. The protocol number for the relevant animals and procedures approved by IACUC is AN200674-00: Title: Role of Growth Factor Signaling in Vascular Physiology” (Approval Date: July 05, 2023). In this study, sex was not considered as a biological variable. Both males and females were used in all experiments.

### MMC-mediated PVOD rat model and administration of C16 and ISRIB

Rats were housed in the vivarium of the cardiovascular research building at UCSF (San Francisco, USA). Both male and female Sprague Dawley young rats (9-10 weeks) and old rats (1.2-1.5 years old) were subjected to following protocols to examine the effect of MMC and/or small molecule inhibitors of ISR (ISRIB; trans-N,N’-(Cyclohexane-1,4-diyl)bis(2-(4-chlorophenoxy)acetamide; IC_50_=5 nM; Sigma-Aldrich, #19785) or PKR (C16; 6,8-Dihydro-8-(1*H*-imidazol-5-ylmethylene)-7*H*-pyrrolo[2,3-*g*]benzothiazol-7-one; IC_50_=210 nM; Sigma-Aldrich, SML0843). MMC was made by dissolving 2 mg MMC in 1 ml saline and was delivered to rats at 3 mg/kg dosage through i.p. injections. Saline was used as vehicle solution for MMC treatment. ISRIB solution was made by dissolving 5 mg ISRIB in 1 ml of dimethyl sulfoxide (DMSO) (Sigma, D2650), followed by dilution to 1 mg/ml and was delivered to rats at 0.25 mg/kg dosage through i.p. injections. The vehicle solution consisted of 1 ml DMSO and 4 ml saline. C16 solution was made by dissolving 10 mg C16 in 1 ml DMSO, followed by dilution to final concentration of 100 μg/ml and was delivered to rats at 33.5 μg/kg dosage through i.p. injections. The vehicle solution consisted of 100 μl DMSO and 10 ml saline.

#### Protocol #1 MMC treatment

Rats were randomly divided into MMC (3 mg/kg) or saline (vehicle)-exposed groups. MMC or saline was administered once by intraperitoneal injection (i.p.) on day 0 (d0). Rats were euthanized on d24 for hemodynamic measurements, RV hypertrophy assessment, and tissue collections.

#### Protocol #2 ISRIB/C16 treatment

Rats were given MMC (3 mg/kg) or vehicle (saline) by i.p. on d0. ISRIB (0.25 mg/kg) or vehicle (DMSO) was given 3 times between d24 and d32 and rats were euthanized on d32. C16 (33.5 μg/kg) or vehicle (DMSO) was given once on d24. On d32, rats were euthanized for hemodynamic measurements, RV hypertrophy assessment, and tissue collections.

### Hemodynamic measurement and tissue histology

The animals were anesthetized with an intraperitoneal injection of a ketamine/xylazine cocktail solution (1 ml ketamine (100 mg/ml) + 100 µl xylazine (20 mg/ml); inject 300 µl per 250 g body weight). A tracheal cannula was then inserted, and the animals were ventilated with room air using VentElite rodent ventilator (Harvard Apparatus) set to maintain respiration at 90 breaths/min and tidal volume at 8 ml/kg body weight. The abdominal and thoracic cavity of the rat was opened carefully to avoid any blood loss, and a 2F pressure-volume catheter (SPR-838, Millar AD Instruments, Houston, TX) was used for RVSP measurements. RVSP was measured while a consistently stabilized pressure wave was shown after the transducer was plugged into RV apex. At the end of the experiments, the hearts and lungs were perfused with phosphate-buffered saline (PBS) for blood removal. Fulton index, or the weight ratio of the right ventricle divided by the sum of left ventricle and septum [RV/(LV + S)], was measured and calculated to determine the extent of right ventricular hypertrophy. Lung, liver, and heart tissues were fixed in 10% formalin for 24 h and then further processed for paraffin sectioning.

The paraffinized lung tissue sections were used for hematoxylin–eosin (H&E)(32) and Gomori’s trichrome staining(33) according to the standard protocol. The images were acquired by Olympus BX51 microscope (Olympus), Ts2 microscopes (Nikon), and Leica SPE confocal microscope. Total areas of fibrotic lesion within each section were quantified using a threshold intensity program from ImageJ.

### Assessment of vascular remodeling

To assess pulmonary artery and vein muscularization, rat lung tissue sections (10 µm in thickness) were subjected to conventional H&E staining. The external and internal diameter of a minimum of 50 transversally cut vessels in tissue block ranging from 25 to 80 µm were measured by determining the distance between the lamina elastica externa and lumen in two perpendicular directions as described previously(34). The vessels were subdivided based on their diameter (microvessels: <50 µm and medium sized vessels: 50-80 µm) and the assessment of muscularization was performed using ImageJ in a blinded fashion by a single researcher to reduce operator variability, who was not aware of the group allocation of the samples being analyzed. The absolute value of the medial thickness was converted to the relative value by setting the medial thickness of vehicle-treated wild type (WT) rats as 1. Additionally, we also assessed the muscularization of pulmonary arteries and veins by the degree of αSMA immunofluorescence staining. The IF signal intensity was quantitated by ImageJ, and the result is presented as a relative signal intensity by setting the value of vehicle-treated WT rats as 1. Images were acquired using Ts2 microscopes (Nikon) and Leica SPE confocal microscope. Airways (bronchi and bronchioles) follow a branching pattern that mirrors the tree-like structure of the lung’s lobes and segments, while veins have a more variable course as they drain blood back to the heart. Airways are typically accompanied by arteries, while veins may be located separately. Furthermore, airways generally have thicker walls compared to veins. When cut in cross-section, are more likely to maintain a round or oval shape, while veins may appear more collapsed or irregular. These characteristics were applied to distinguish veins from airways.

### Immunoblot analysis

The rat tissue lysates were prepared in the lysis buffer (1% Triton X-100, 150mM NaCl, 50mM Tris-Cl at pH 7.5, 1mM EDTA). The supernatants were collected, and total protein concentration was measured by NanoDrop 2000c (Thermo Scientific). Proteins samples were denatured in SDS-sample buffer for 5 min at 95°C and loaded onto Mini-Protean TGX^TM^ gels (BioRad Laboratories) in equal amounts and subjected to electrophoresis. Nitrocellulose membrane (Genesee Scientific) was used to blot the gels, which were blocked with 5% non-fat milk or 3% BSA in 1x Tris buffered saline with 0.1% tween-20 (1x TBST) for 1 h at RT. The membranes were incubated at 4 °C overnight with a primary antibody. Chemiluminescence signals were detected using SuperSignal™ West Dura extended duration substrate (ThermoFisher) and imaged using an Odyssey Dlx Imaging System (LI-COR). Antibodies used for immunoblots are found in Supplementary Information. The quantity of each protein was normalized to the amount of a loading control protein. Subsequently, its relative quantity was calculated by setting the amount of the protein in the control (vehicle-treated) sample as 1.

### Immunoprecipitation assay

Rat tissue and plasma samples were lysed in IP buffer (1% Triton X-100, 150mM NaCl, 50mM Tris-Cl at pH 7.5, 1mM EDTA) supplemented with protease inhibitors (1:100 dilution) and phosphatase inhibitor (1:100 dilution). Lysates were nutated for 30 min at 4 °C, followed by centrifugation at 12,000 g for 10 min and supernatant were collected. One tenth of the lysate was saved as an input sample for immunoblot. The lysate was incubated with indicated antibodies and anti-IgG (negative control) nutating overnight at 4°C followed by the addition of dynabeads^TM^ Protein A/G and rocking for 4 h at 4 °C. The magnetic beads were precipitated and rinsed thrice with IP buffer for 5 min at 4 °C. The washed elute was boiled at 95°C for 8 min in a sample loading buffer and subjected to immunoblot along with input. For the input samples: initially, the quantity of the indicated protein was normalized to the amount of a loading control protein, such as beta-actin. Subsequently, its relative quantity was calculated by setting the amount of the protein in the vehicle-treated sample as 1. For the IP samples, the amount of the indicated protein in MMC-treated sample was presented with the protein amount in the control (vehicle-treated) sample set as 1.

### Reverse Transcriptase-quantitative Polymerase Chain Reaction (qRT-PCR)

Total RNA was isolated from Rat lung tissue and subjected to cDNA preparation by the reverse transcription reaction using an iScript cDNA Synthesis Kit (#17088890, Bio-Rad). qPCR analysis was performed in triplicate using iQ SYBR Green Supermix (#1708882, Bio-Rad). The relative expression values were determined by normalization to *GAPDH* transcript levels and calculated using the ΔΔCT method. qRT-PCR primer sequences are found in Supplementary Information.

### Immunofluorescence staining

Anesthetized rats were flushed with 1X PBS and fixed in 4% paraformaldehyde (w/v), transferred to 1X PBS after 24 h, and subjected to paraffin embedding. Additionally, the right bronchus of flushed lungs was sutured, and the left lung inflated with 1% low melt agarose for fixation and paraffin embedding, and the right lung split for snap freezing for protein and RNA studies.

Immunofluorescence was performed using antibodies listed in the Supplementary information. For SMA and EC staining, sections were subjected to deparaffinization, antigen retrieval, and permeabilization followed by blocking and primary antibody incubation overnight at 4°C. A solution of Alexa Fluor secondary antibodies (Invitrogen) was applied for 2 hours at RT. IF images were acquired using a confocal microscope (Leica SPE) or Eclipse Ts2 Inverted LED phase contrast microscope (Nikon) and analyzed using ImageJ. Antibodies used are found in the Supplementary Information.

### Statistical analysis

All numerical data are presented as mean ± standard error of the mean (SEM). Statistical analysis was performed using Microsoft Excel and GraphPad Prism 10 (La Jolla, CA). Datasets with two groups were subjected Student’s *t*-test, unpaired, equal variance whereas comparison among three or more than three groups was done by ANOVA followed by Tukey’s post hoc corrections. Analysis of variance was applied to experiments with multiple parameters, one-or two-way as appropriate. And, where required, significance was analyzed using a post hoc Tukey test and indicated as P-values.

## Authors Contributions

The authors confirm contribution to the paper as follows. *Study conception and design*: Amit Prabhakar and Akiko Hata. *Execution of experiments*: Amit Prabhakar, Meetu Wadhwa, Rahul Kumar, Prajakta Ghatpande,. *Data analysis*: Amit Prabhakar, Meetu Wadhwa, Rahul Kumar, Giorgio Lagna, and Akiko Hata. *Interpretation of results*: Amit Prabhakar, Meetu Wadhwa, Rahul Kumar, Brian Graham, Giorgio Lagna, and Akiko Hata. *Primary draft manuscript preparation*: Amit Prabhakar and Akiko Hata. All authors reviewed the results and approved the final version of the manuscript.

## Acknowledgements

Microscopy data were acquired at the Center for Advanced Light Microscopy-CVRI Microscopy core on microscopes. We thank Ms. Ananyaa Arvind (UCB), Ms. Simren Gupta (UCSB), Mr. Sehaj Dhami (Santa Clara Univ.), and Ms. Yuhang Yao (Tshinghua Univ.) for technical support. Grant funding was provided by National Heart, Lung, and Blood Institute (NHLBI; R01HL132058, R01HL153915, and R01HL164581) to A.H.; American Heart Association 19CDA34730030, Cardiovascular Medical Research Fund (CMREF) and United Therapeutics Jenesis Innovative Research Award to R.K.; by NHLBI (R01HL135872 and P01HL152961) to B.B.G.s

---

**Supplementary Fig. S1.**
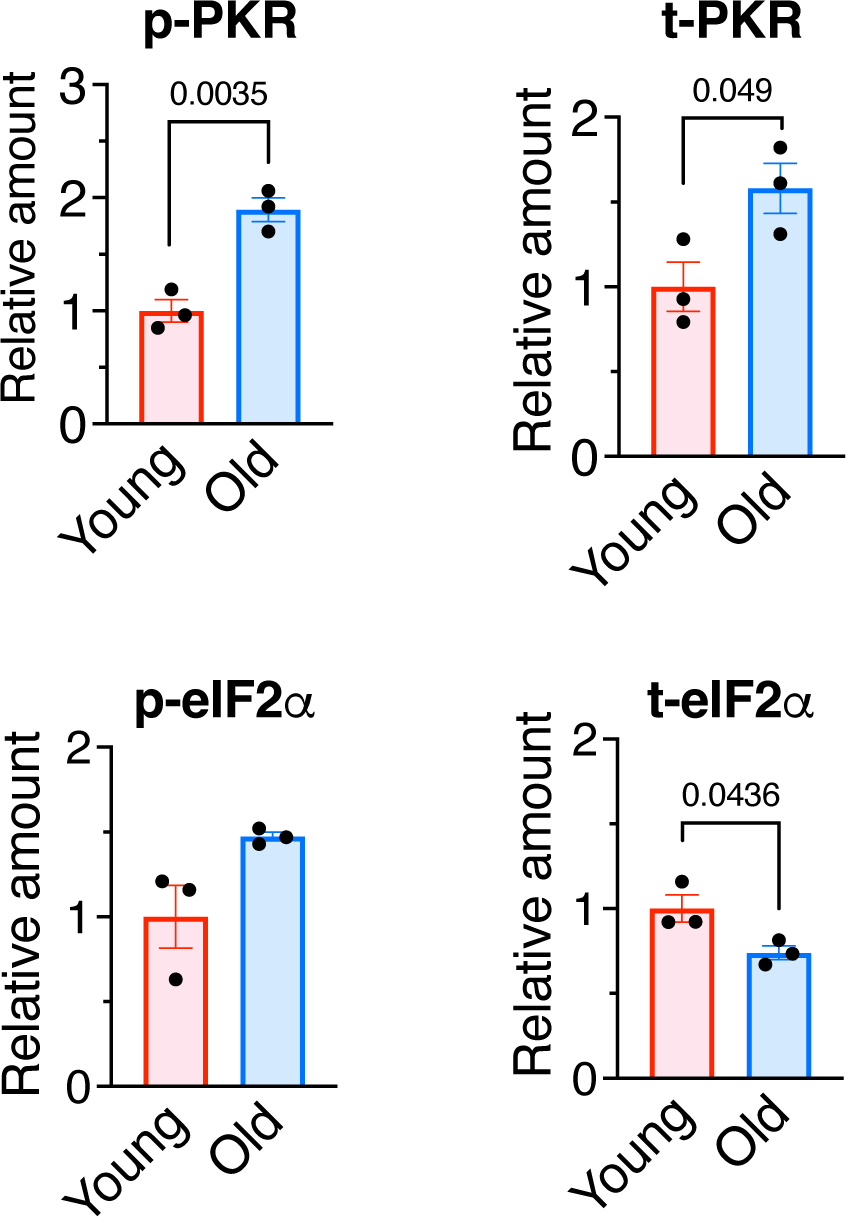
Differential ISR activity in young and old rats a. Quantitation of immunoblot of p-PKR, t-PKR, p-eIF2α, and t-eIF2α in the lung lysates of young and old rats. The amount of indicated proteins normalized to β-actin and shown as mean±SEM (right). n=3 independent samples per condition. Statistical analysis was performed using two-tailed Student’s t-test with p<0.05.

